# Transcriptome analysis in *C. elegans* early embryos upon depletion of the Topoisomerase 2/condensin II axis

**DOI:** 10.1101/2024.10.17.618010

**Authors:** Mezmur D. Belew, W. Matthew Michael

## Abstract

In *C. elegans*, the chromosome compaction factors topoisomerase 2 (TOP-2) and condensin II have been shown to globally repress transcription in multiple contexts. Our group has previously reported that TOP-2 and condensin II repress transcription in the *C. elegans* germline during larval starvation, oocyte maturation, and in germline progenitor cells of the early embryo. Here, we assess the transcriptome of early embryos treated with RNAi against TOP-2 and the condensin II subunit CAPG-2. We found 144 upregulated and 172 downregulated genes. Further analysis showed that the upregulated genes are mostly somatic, with a host of neuronal cells present in our tissue enrichment analysis.

## Introduction

The role chromatin organization plays in regulating gene transcription has been reported in multiple organisms and cellular contexts. The chromatin modifiers topoisomerase 2 (TOP-2) and condensins have been shown to modulate gene transcription in organisms ranging from yeast to humans (Herrero-Ruiz et al., 2021; Nikolaou et al., 2013; Bhalla et al., 2002; Albritton and Ercan, 2018). For instance, in budding yeast, TOP2 has been reported to act on the promoter regions of select genes, and loss of TOP2 activity results in the derepression of a large number of genes (Pedersen et al., 2012). Similarly, in quiescent yeast cells, condensin facilitates long-range interactions that mediate genome-wide transcriptional silencing (Swygert et al., 2019). In *C. elegans*, our group has shown that TOP-2 and condensin II work together to repress transcription at multiple time points throughout the development of the germline. During larval starvation (L1 diapause), TOP-2 and condensin II work with heterochromatin pathway proteins to form the global chromatin compaction (GCC) pathway (Belew et al., 2021). This pathway condenses chromatin and suspends transcription in the primordial germ cells (PGCs) (Belew et al., 2021). In addition, components of the GCC pathway, along with PIE-1 (a transcriptional repressor shown to be active in early embryos), also repress transcription in maturing oocytes (Belew et al., 2023). Lastly, we have recently shown that TOP-2 and condensin II also repress transcription in P blastomeres of the early embryo (Belew et al, 2024). This work follows up on the early embryo study by looking at how loss of TOP-2 and condensin II impacts the early embryonic transcriptome.

## Materials and Methods Animal growth

The N2 (wild-type) strain was used in this study. Worms were maintained on 60mm plates containing nematode growth media (NGM) seeded with the *E. coli* strain OP50 or HT115. Worms were grown at 20° C and propagated through egg preparation (bleaching) every 72 hours.

### Bacterial strains

OP50 bacteria served as the primary food source. It was grown in LB media containing 100 ug/ml streptomycin by shaking at 37° C overnight. 500 ul of the culture was seeded on Petri-dishes containing NGM + streptomycin. HT115 bacteria grown in LB media containing 100 ug/ml carbenicillin and 12.5 ug/ml tetracycline and seeded on NGM + carbenicillin + tetracycline plates were also used as a source of food. Our RNAi strains were obtained from the Ahringer library and verified by Sanger sequencing. Bacteria containing dsRNA were streaked on LB-agar plates containing 100 ug/ml carbenicillin and 12.5 ug/ml tetracycline and incubated at 37° C overnight. Single colonies were then picked and grown in 25ml LB cultures with 100 ug/ml carbenicillin and 12.5 ug/ml tetracycline. 500 ul of this culture was seeded on 60 mm Petri-dishes containing 5mM IPTG.

### Egg preparation

Bleach solution containing 3.675 ml H2O, 1.2 NaOCl, and 0.125 ml 10N NaOH was prepared. Adult worms were washed from plates with 5 ml of M9 minimal medium (22mM KH2PO4, 22mM Na2HPO4, 85mM NaCl, and 2mM MgSO4). Worms were centrifuged at 1.9 KRPM for 1 minute and the excess medium was removed, then the bleach solution was added. Eggs were extracted by vortexing for 30 seconds and shaking for 1 minute. This was done a total of 3 times and worms were vortexed one last time. Then the eggs were spun down at 1900 rpm for 1 minute and excess bleach solution was removed and the eggs were washed 3 times with M9 minimal medium.

### RNAi treatment

RNAi containing NGM plates were prepared as described in the “Bacterial strains” section. For double RNAi treatments, RNAi cultures were mixed at a 1:1 ratio by volume. HT115 cells transformed with an empty pL4440 vector was used as a negative control. RNAi conditions used in this study and tests for their efficacy is described below:*top-2/capg-2* double RNAi

Worms were grown on HT115 food plates for the first 24 hours and were transferred to *top-2/capg-2* double RNAi plates for the next 48 hours. Embryonic lethality ranged from 90%-100% for this RNAi treatment.

### RNA isolation for sequencing

Embryos were synchronized by bleaching young adults as soon as the first eggs were seen in the animal to enrich for early embryos (Boeck et al., 2016). This was done on three successive generations to tighten the synchrony. After the final round of bleaching, the embryos were washed with 10 ml cold M9 minimal medium and packed by centrifuging at 1.9 KRPM. A portion of the sample was taken to assess the stage of the embryos by staining DNA. The rest of the embryos were flash-frozen in liquid nitrogen and stored at -80oC until the RNA isolation step.

Samples were thawed and four volumes of Tri-reagent (Sigma Aldrich, St. Louis, Missouri) were added to them. The mixture was then vortexed vigorously to solubilize the embryos. Once solubilized, the sample was distributed into Eppendorf tubes by adding 1250 ul of the sample to each tube. Samples were incubated for 5 minutes at room temperature and centrifuged at 14 KRPM for 10 minutes at 4oC to remove debris. The supernatant was then transferred into fresh tubes and 200 ul of Chloroform (Sigma Aldrich, St. Louis, Missouri) was added to each tube. The solution was vortexed for 15 seconds and incubated at room temperature for 3 minutes before being centrifuged at 14 KRPM and 4oC for 15 minutes. The top aqueous layer was carefully transferred to new centrifuge tubes and 500 ul of isopropanol (Sigma Aldrich, St. Louis, Missouri) was added to it. This mixture was incubated at room temperature for 10 minutes and then centrifuged for 10 minutes at 14 KRPM and 4oC to precipitate RNA. The supernatant was discarded, and the pellet was washed with 100 ul of 75% EtOH in diethyl pyrocarbonate (DEPC) treated water. The sample was then centrifuged one more time at 7.5 KRPM for 5 minutes at 4oC, the supernatant was removed, and the pellet was air-dried in a fume hood for 15 minutes. Finally, the dry pellet was dissolved in 50 ul of DEPC treated water and sent to Genewiz (South Plainfield, NJ) for library preparation and sequencing. This was repeated for a second biological replicate.

### RNA sequencing data analysis

Library preparation and sequencing were done by Genewiz (South Plainfield, NJ). Fastq files were imported into Partek-flow. Prior to alignment, reads were assessed for sequencing quality and bases with quality score less than 20 were trimmed. If a read had a length lower than 25 bp after trimming the whole read was discarded. STAR aligner was used to align reads to *C. elegans* – WB235 assembly which was retrieved from WormBase. After alignment, reads were quantified using the Partek-EM method to WS235manual-Ensembl104 annotation model that is also available on WormBase. Genes whose counts across all samples were less than 50 were filtered out and then gene counts were normalized using the median ratio method for DEseq2. Finally, DEseq2 was used to generate a list of differentially expressed genes.

Upon getting the list of differentially expressed genes, we set an adjusted P-value cutoff of 0.05 to attribute statistical significance. A fold-change cutoff of -1 and 1 was used to identify genes that were downregulated and upregulated, respectively. The “ggplot2” package was on R used to construct a volcano plot to visualize these genes. Principal component analysis to see the separation of our samples was done using R and was visualized using the “ggplot2” package. The “Gene set enrichment analysis” tool on WormBase was used to identify tissues and biological processes that were enriched in our upregulated genes. A short R script that cross-checks our list of upregulated genes against lists of genes from various studies for similarities was written to perform the overlap analyses presented in this study.

### HCR (In Situ Hybridization Chain Reaction)

A kit containing a DNA probe set, DNA hybridized chain reaction (HCR) amplifier hairpins, and hybridization, wash, and amplification buffers were purchased from Molecular Instruments (molecularinstruments.com). Genes that were examined were *vet-6*, F58E6.6, *ppw-2, cab-1, zip-2*, and *asns-2*. DNA was visualized with Hoechst 33342 dye. Embryos were prepared by bleaching and were immediately spotted on poly-L-lysine coated slides. Coverslips were applied and slides were freeze-cracked to permeabilize samples. Immediately after freeze-cracking, 500 ul of 100% ice-cold methanol was applied over the samples. Slides were dried off by tilting slides, 1 ml of 4% paraformaldehyde (PFA) was added and samples were incubated in a humidity chamber for 10 minutes. Samples were then washed 3 times with 100 ul of PBS-T. A 1:1 solution of probe hybridization buffer (PHB) and PBS-T was added to the samples, and they were incubated for 5 minutes at room temperature. Samples were then prehybridized with PHB for 30 minutes at 37oC and DNA probes (at a final concentration of 2 picomoles per 500 ul of PHB) were added to the samples and were incubated overnight at 37oC.

The next day, samples were washed 4 times with probe wash buffer (PWB) at 37oC with 15 minutes of incubation for each wash. They were then washed three more times with 5xSSCT at room temperature. Samples were pre-amplified with Amplification buffer for 30 minutes at room temperature. Probe amplifier hairpins were snap cooled by heating to 95oC for 90 seconds and putting in a dark drawer for 30 mins then were added to the sample. Worms were incubated with the hairpins overnight in a dark drawer.

On the third day, samples were washed with 5xSSCT and incubated with Hoechst-33342 (1:5000 dilution) for 15 minutes. Finally, samples were mounted on poly-L-lysine coated slides and imaged.

### Sample imaging

All slides were imaged using an Olympus Fluoview FV1000 confocal microscope using Fluoview Viewer software. Laser intensity was controlled for experiments to achieve consistency among samples.

### Reagents

N2 strains were provided by the CGC, which is funded by NIH Office of Research Infrastructure Programs (P40 OD010440).

## Results and Discussion

Recent work has shown that loss of TOP-2/condensin II allows precocious transcription in P blastomeres of the early embryo (Belew et al, 2024). In this work we were interested in determining how loss of these factors impacts the transcriptiome in early embryos. For this, we performed RNA-seq analysis on early embryos treated with either control RNAi or RNAi against TOP-2 and the condensin II subunit CAPG-2 (*top-2*/*capg-2* RNAi). It is well established that condensins and topoisomerase 2 work together to promote chromosome compaction in preparation for mitosis (Shintomi et al., 2015; Kinoshita and Hirano, 2017), and previous work from our group showed that co-depletion of TOP-2 and CAPG-2 obstructed genome silencing during oocyte maturation (Belew et al., 2023), and thus, we used the same RNAi conditions here to study how co-depletion would impact gene expression in early embryos. L1 larvae were synchronized by hatching in minimal media and then plated on food. Growth was monitored such that as soon as young adults had formed embryos the samples were bleached and early embryos were purified and used for RNA extraction. The transcriptomes were sequenced in replicate, and principal component analysis revealed that replicates of the *top-2*/*capg-2* RNAi treated samples cluster away from those of the control RNAi treated samples, confirming RNAi treatment underlies the separation among the samples (Figure 1). Moreover, using an adjusted p-value cutoff of 0.05, we found 316 differentially expressed genes (DEGs) in the *top-2/capg-2* RNAi samples, with 144 of them upregulated and 172 downregulated (Figure 2, Extended data 1). The fact that more than half of the DEGs are downregulated was surprising, given the repressive nature of TOP-2 and condensin II (Belew et al., 2021; 2023; 2024; Swygert et al., 2019). A possible cause for this could be the misexpression of repressor genes due to the loss of TOP-2 and condensin II which, in turn, would result in the downregulation of the genes in our experiment. A deeper analysis of the DEGs revealed that they were evenly distributed among the six *C. elegans* chromosomes, implying the occurrence of genome-wide misexpression without a bias towards a single chromosome.

**Figure 1.**
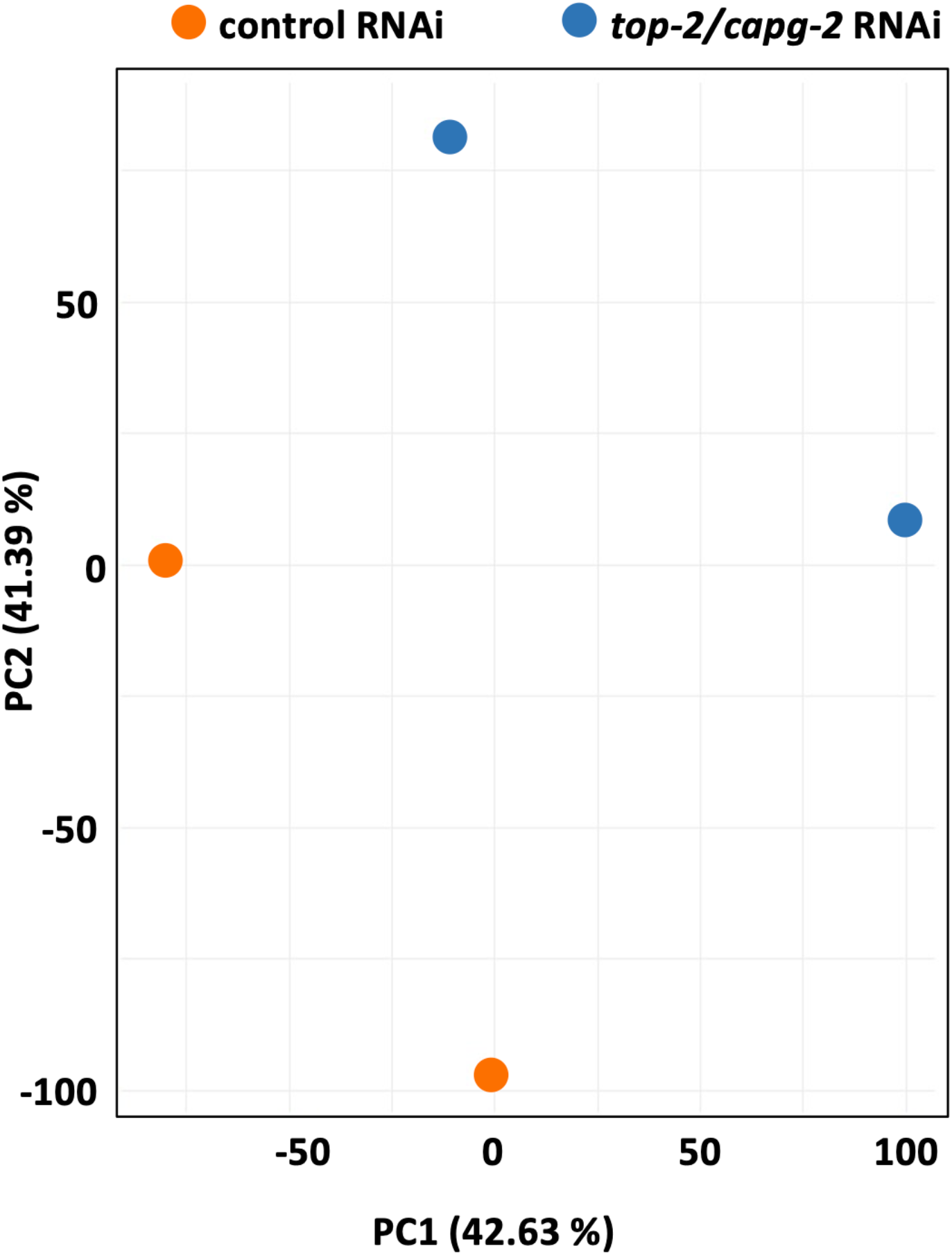
Principal component analysis shows that separation of the data sets is linked to the RNAi that the animals were exposed to.

**Figure 2.**
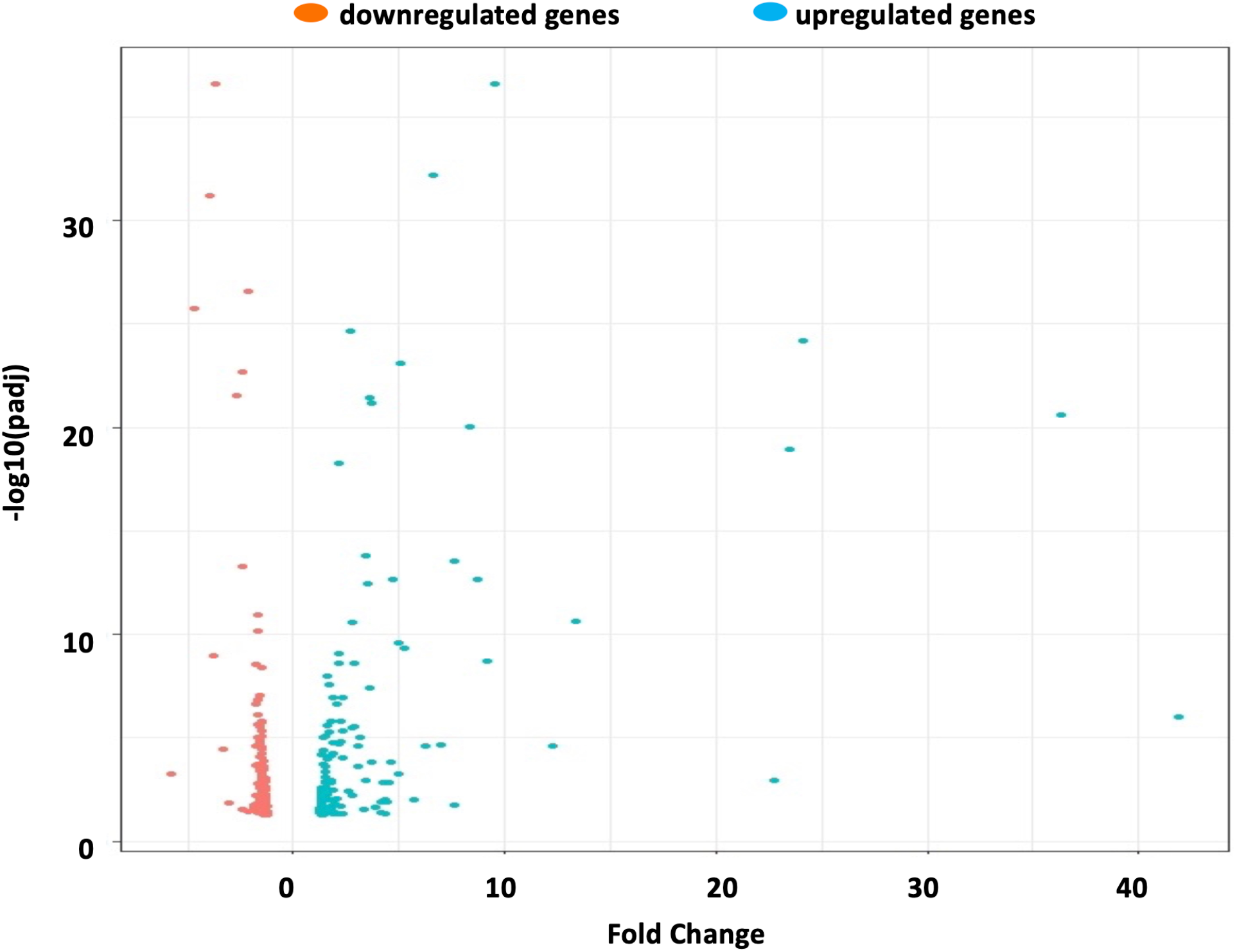
A volcano plot for the differentially expressed genes. The x-axis shows fold-change and the y-axis shows plots adjusted P-values (see Methods).

We next analyzed our group of 144 upregulated genes to search for commonality. Gene enrichment analysis was performed, and we observed that many of the genes (22 in total) shared gene ontology (GO) terms related to defense and interspecies interactions (Figure 3A). For tissue distribution, most of the enrichment we observed was for the nervous system, as SIB, ALA, DVB, and SIA are all neurons (Figure 3B). Next, we compared our dataset to other previously published datasets to get more information regarding our upregulated genes. First, because TOP-2 and condensin II work together in the GCC pathway with the SET-25 and MET-2 methyltransferases (Belew et al., 2021), we compared our list of upregulated genes to a list of genes that are upregulated in *set-25, met-2* double mutant early embryos, produced by Gasser and colleagues (Zeller et al, 2016). The two groups had just 5 genes in common (Table 1), suggesting that TOP-2/condensin II acts independently of the H3K9me pathway in early embryos. Similarly, we observed scant overlap with transcripts that are provided maternally to early embryos (Figure 1E; Quarato et al., 2021), or with genes that are expressed in the gonad (Table 1; Reed et al., 2020). By contrast, strong overlap was observed with genes that are expressed somatically (Table 1; Reed et al., 2020). Taken together, these analyses indicate that the majority of the upregulated genes are somatic genes that are not maternally supplied in normal conditions.

**Figure 3.**
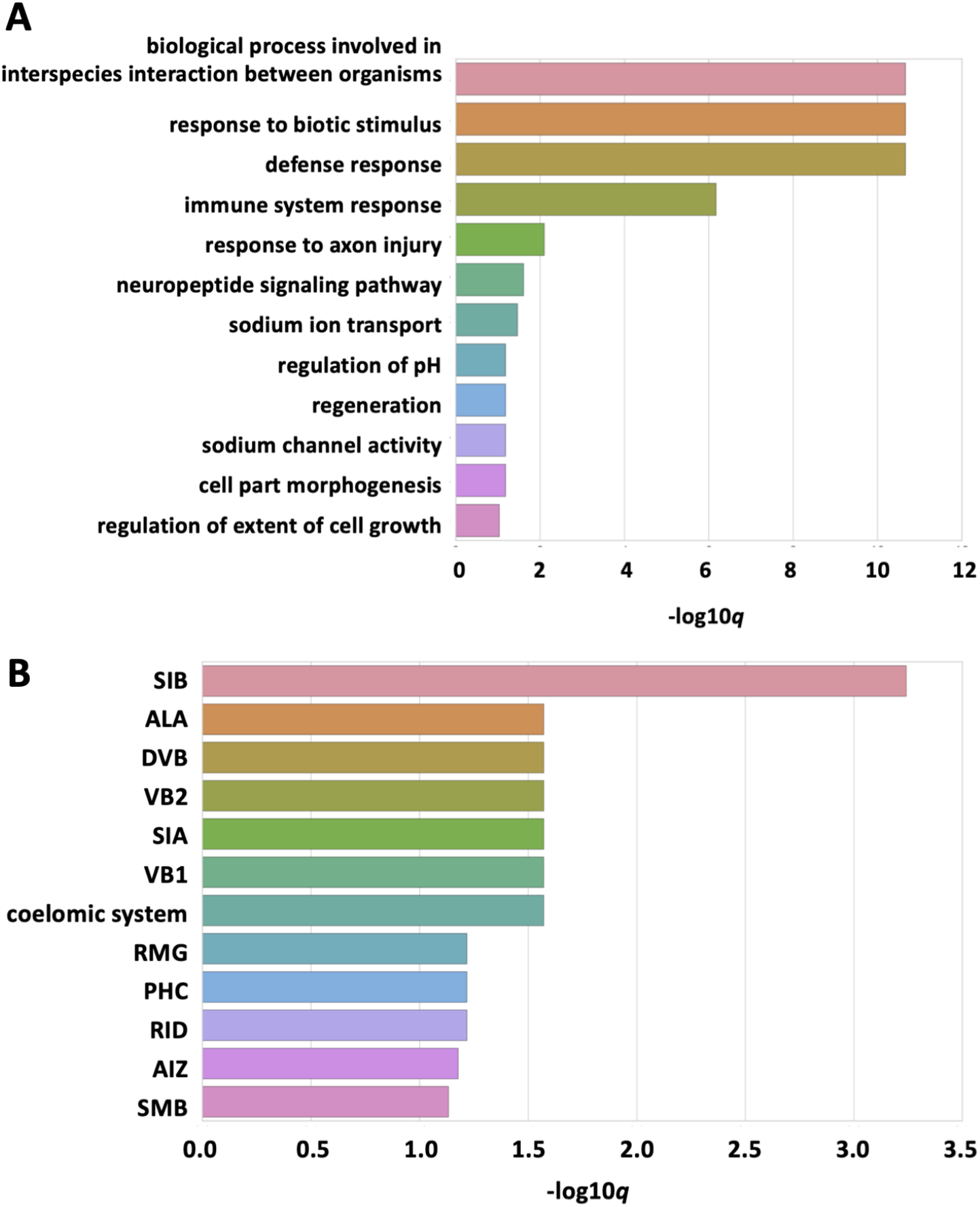
A) Gene ontology (GO) analysis for our set of 144 upregulated genes were performed using the Gene Set Enrichment Analysis tool that is available on Wormbase.org (https://wormbase.org/tools/enrichment/tea/tea.cgi). B) Tissue enrichment analysis for our set of 144 upregulated genes were performed using the same tool as in (A).

**Table 1.** Overlap analysis for our set of 144 upregulated genes was performed against the indicated data sets.

| Condition | # of genes | # of the 144 genes upregulated<br>after *top-2/capg-2* RNAi<br>found in data set | Reference |
| --- | --- | --- | --- |
| Genes upregulated in *met-2/set-25* double mutants | 430 | 5 (3.5%) | Zeller et al., 2016 |
| Maternally expressed genes | 7088 | 28 (19.4%) | Quarato et al., 2021 |
| Gonad-enriched genes | 3207 | 5 (3.5%) | Reed et al., 2020 |
| Soma-enriched genes | 8055 | 106 (73.6%) | Reed et al., 2021 |

Lastly, we wanted to validate our findings from the RNA sequencing experiment using a second set of independent experiments. To do so, we picked three genes that are upregulated in our *top-2/capg-2* RNAi samples (*zip-2, cab-1*, and *asns-2*) and performed *in situ* HCR to assess their expression patterns in early embryos by directly visualizing their transcripts (Choi et al., 2018). We looked at 4-and 8-cell embryos and found that wild-type embryos had little to no HCR signal. However, upon the depletion of TOP-2 and CAPG-2, we saw a pan-embryonic increase in transcripts for all three genes (Figures 4&5), corroborating the list of differentially expressed genes in our experiment.

**Figure 4.**
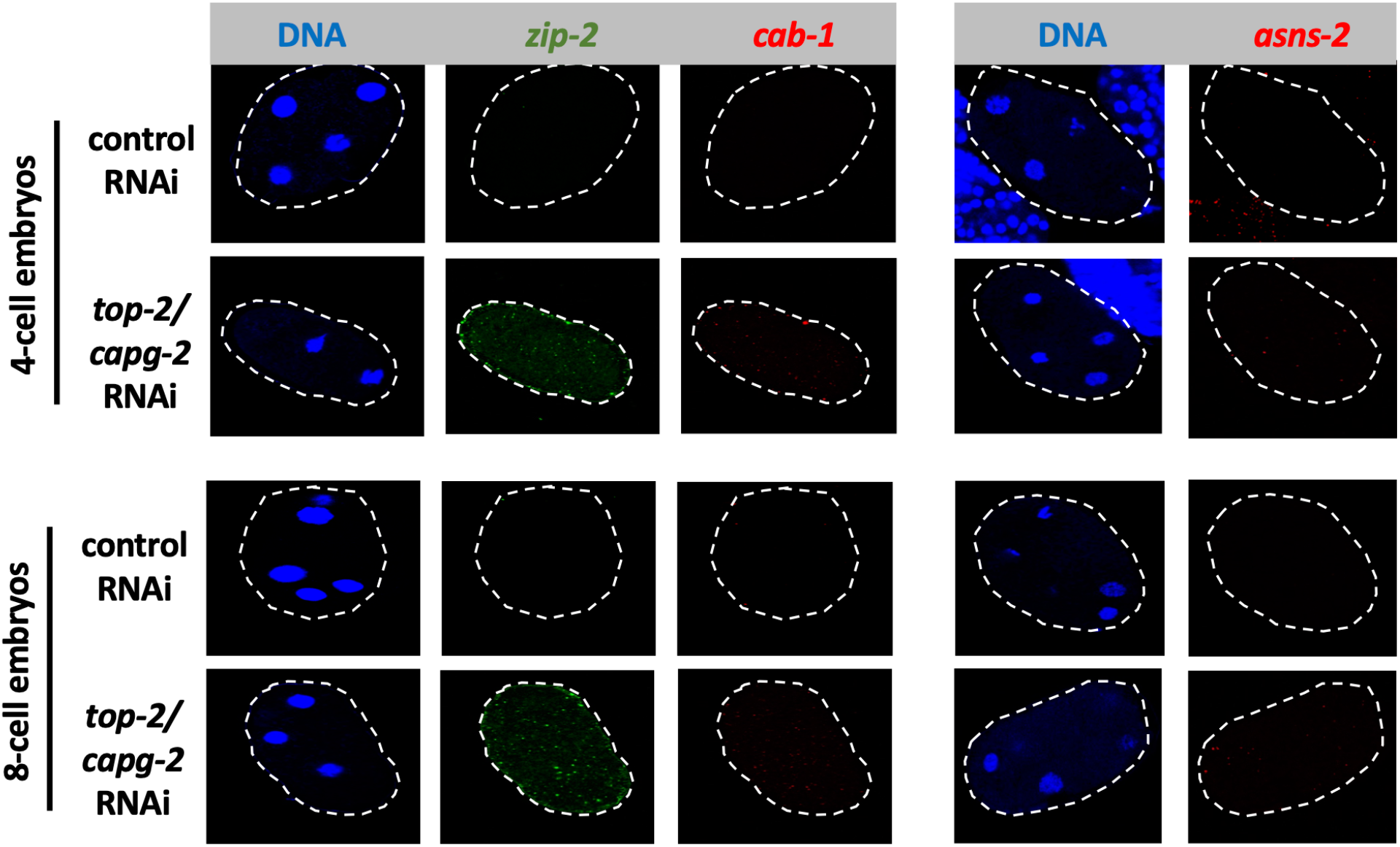
Samples were treated with either control or *top-2/capg-2* RNAi. HCR was performed to probe for *zip-2* transcripts (green), *cab-1* transcripts (red), *asns-2* transcripts (red) and DNA (blue) in 4- and 8-cell embryos. Representative images are shown.

**Figure 5.**
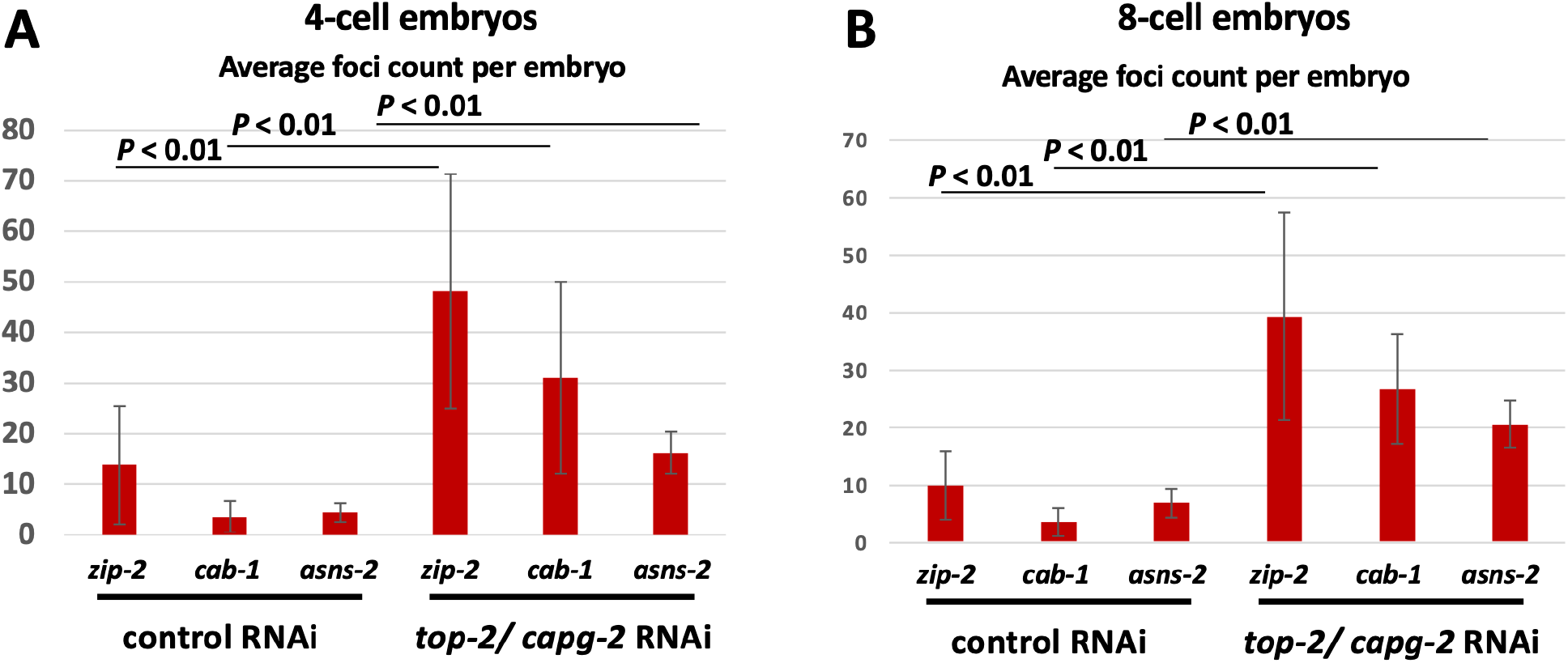
Quantification of the HCR experiment shown in Figure 4. HCR foci were counted for the indicated samples and their averages are plotted and compared to one another using Student’s t-test.

## Author Contributions

Mezmur Belew: Conceptualization, Formal analysis, Investigation, Methodology, Visualization, Writing - original draft.

W. Matthew Michael: Conceptualization, Funding acquisition, Project Administration, Visualization, Writing – review & editing.

## Acknowledgements

We are very grateful to Meng Li and Yibu Chen of the USC Libraries Bioinformatics Service Program for help with data analysis.

## Tables and Figures

**Extended data 1**: A table containing the complete list of differentially expressed genes from our RNA seq analysis of early embryos treated with either control or *top-2/capg-2* RNAi.

